# Targeted Reactivation of X-linked Endogenous FOXP3 Gene from X-chromosome Inactivation in Human Female Breast Cancer Cells

**DOI:** 10.1101/2021.08.14.456310

**Authors:** Xuelian Cui, Zhifang Xu, Shuaibin Wang, Xin Li, Erica Stringer-Reasor, Sejong Bae, Leiping Zeng, Dehua Zhao, Runhua Liu, Lei S. Qi, Lizhong Wang

## Abstract

Unlike autosomal tumor suppressors, which must be inactivated by a two-hit Knudson mechanism, X-linked tumor suppressors can be inactivated by a single hit due to X-chromosome inactivation (XCI). Here, we argue that targeted reactivation of the non-mutated allele from XCI offers a potential therapy for female breast cancers. Towards this goal, we developed a dual CRISPR interference and activation (CRISPRi/a) approach for simultaneously silencing and reactivating multiple X-linked genes using two orthogonal, nuclease-deficient CRISPR/Cas9 (dCas9) proteins. We verified the efficacy for use of *Streptococcus pyogenes* (Sp) dCas9-KRAB for silencing *XIST* and *Staphylococcus aureus* (Sa) dCas9-VPR for activating *FOXP3* in various cell lines of human female breast cancers. We confirmed CRISPR reactivation of the non-mutated copy of *FOXP3* from XCI in human breast cancer CRL2316 cells, which express a synonymous heterozygous mutation (p.L266L; c.798G>C) in the coding region of *FOXP3*. Further, simultaneous silencing of *XIST* from XCI led to enhanced and prolonged *FOXP3* reactivation. We optimized CRISPRa by fusing SadCas9 to the demethylase TET1 and observed enhanced *FOXP3* activation. Analysis of the conserved CpG-rich region of *FOXP3* intron 1 confirmed that CRISPRi/a-mediated simultaneous *FOXP3* activation and *XIST* silencing were accompanied by elevated H4 acetylation, including H4K5ac, H4K8ac, and H4K16ac, and H3K4me3 and lower DNA methylation. This indicates that CRISPRi/a targeting to *XIST* and *FOXP3* loci alters their transcription and their nearby epigenetic modifications. The simultaneous activation and repression of the X-linked, endogenous *FOXP3* and *XIST* from XCI offers a useful research tool and a potential therapeutic for female breast cancers. This approach also provides new routes of targeted therapy for other X-chromosome-linked genetic disorders.

## Background

Females have a unique mechanism of X-chromosome inactivation (XCI), which silences X-linked genes from one of the two alleles (1, 2), leading to a dose compensation of X-linked genes between sexes. Mechanistically, silenced chromatin in XCI is coated by the non-coding RNA X-inactive specific transcript (*XIST*), potentially via *XIST* binding to X-linked gene loci for XCI initiation (3, 4). XCI, as a paradigm for large-scale epigenetic regulation (5–7), is triggered by the polycomb repressive complex 1/2 (PRC1/2)- and *XIST*-binding proteins (8–12). The PRC1/2- and *XIST*-binding proteins initiate the accumulation of non-coding RNA *XIST* on the inactivated X chromosome, followed by a series of epigenetic modifications (13, 14). However, XCI is reversible, and, in human female cells, at least 23% of X-linked genes escape XCI (1,14–18). In XCI, repressive histone modifications with DNA methylation prevent transcription of X-linked genes, whereas permissive histone modifications, DNA hypomethylation, and RNA polymerase II are enriched at escape loci for transcription of X-linked genes from XCI (19). X-Chromosome reactivation can occur during early embryogenesis; in various stem cells; during the reprogramming of somatic cells to pluripotent stem cells; and upon combined *XIST* inhibition, DNA demethylation, and histone deacetylation (14, 20). However, restoring all X-linked gene expression in the XCI will likely bring about undesired results since the resulting elevated levels of X-linked genes can be toxic for female cells.

In certain tumors, autosomal tumor suppressor genes can be inactivated by a two-hit mechanism (both alleles are inactivated, known as the Knudson hypothesis). However, X chromosome-linked tumor suppressor genes, such as *FOXP3* at Xp11.23 (21), *WTX* at Xq11.2 (22), and *ATRX* at Xq21.1 (23, 24), can be inactivated by a single-hit mechanism (only one allele is inactivated), because XCI inactivates the other allele. In female breast cancer cells, all identified gene deletions of *FOXP3* are heterozygous, and mice with a *Foxp3*-heterozygous mutation develop spontaneous breast cancers, suggesting that the active allele may be the only allele affected (21). Similarly, all identified mutations and deletions of *WTX* for female patients with Wilms’ tumors are heterozygous (22). These data indicate that, for females with cancer, it may be possible to reactivate the non-mutated, inactivated allele for therapeutic purposes. Recent research has shown that X-linked genes can be reactivated from XCI by targeted editing of DNA methylation (25), offering a possibility for targeted reactivation of X-linked genes from XCI.

Recent advances in genome engineering led to the development of endonuclease-deficient CRISPR/Cas9 protein (dCas9) for dynamic, tunable, and reversible transcriptional and epigenome regulation of endogenous genes (26, 27). We recently developed, for complex gene regulation, a flexible CRISPR/dCas9-based platform that independently controls the expression of various genes (repression and activation) within the same cell (28). Thus, for therapeutic purposes, targeted reactivation of XCI-endogenous tumor suppressor genes may be an effective strategy to restore their function in female cancer cells. The X-linked FOXP3 gene has dual roles in tumor cells and immune cells. As a master transcriptional regulator of regulatory T cells (Tregs), FOXP3 limits antitumor immunity (29), whereas, in breast cancer cells, it is an epithelial cell-intrinsic tumor suppressor (21,30–36) and is implicated in a tumor-suppressive function in the inhibition of tumor initiation and progression (34,35,37–41). Thus, using *FOXP3* as an X-linked model gene, we aimed to develop, for human female breast cancer cells, a tunable and reversible, targeted reactivation of X-linked tumor suppressor genes. In the present study, using CRISPR interference and activation (CRISPRi/a), we achieved, for human female breast cancer cells, targeted reactivation of X-linked endogenous *FOXP3*, at least a partial reactivation from XCI. Next, we investigated the potential epigenetic mechanism during CRISPRi/a-mediated reactivation of *FOXP3*.

## Results

### Establishment of a flexible CRISPRi/a-based platform for targeted repression of *XIST* and activation of *FOXP3*

dCas9-mediated, gene-specific epigenetic modifications are based on dCas9 fusion to transcriptional and epigenetic effectors (42). For example, dCas9 fusion to the transcriptional repressor Krüppel-associated box (KRAB) domain (43) is used for CRISPRi-mediated repression of gene expression. In contrast, dCas9 fusion to tripartite transcriptional activators, including herpesvirus protein VP64, NF-κB p65 (activation domain), and Epstein–Barr virus R (Rta) (44, 45) (VPR) is used for CRISPRa-mediated activation of gene expression. To induce the transcription of X-linked endogenous *FOXP3* in female cells, we developed a dCas9-based platform for simultaneous *XIST* repression and *FOXP3* activation within the same cells using our previously established method (28). As shown in Supplementary Table S1, DNA constructs consist of a coherent set of pSLQ1922 for *Streptococcus pyogenes* (Sp) dCas9 (SpdCas9)-KRAB [doxycycline (Dox) inducible], pSLQ1932 for SpdCas9-VPR (Dox inducible), pSLQ2840 for *Staphylococcus aureus* (Sa) dCas9 (SadCas9)-VPR, modified pSLQ2840 for SadCas9-TET1, pSLQ2837 for *XIST* single guide RNA (sgRNA), and pSLQ2806 for *FOXP3* sgRNA. The three *XIST* sgRNA1/2/3 were established in our previous study (46) (Fig. 1A and Supplementary Table S2). For targeted activation of *FOXP3,* we designed five sgRNAs (Supplementary Table S2) at the two CpG-rich loci in the proximal promoter region of the transcription start site and the conserved non-coding sequence (CNS) region of intron 1 (47–50) of *FOXP3* (Fig. 1B). The *XIST-* and *FOXP3-* sgRNAs guide the SpdCas9-KRAB and SadCas9-VPR fusions, respectively, to the target sequences at the *XIST* (proximal promoter region) and *FOXP3* (two CpG sites) loci, respectively (Figs. 1A and 1B) (28,46,50–57).

**Figure 1.**
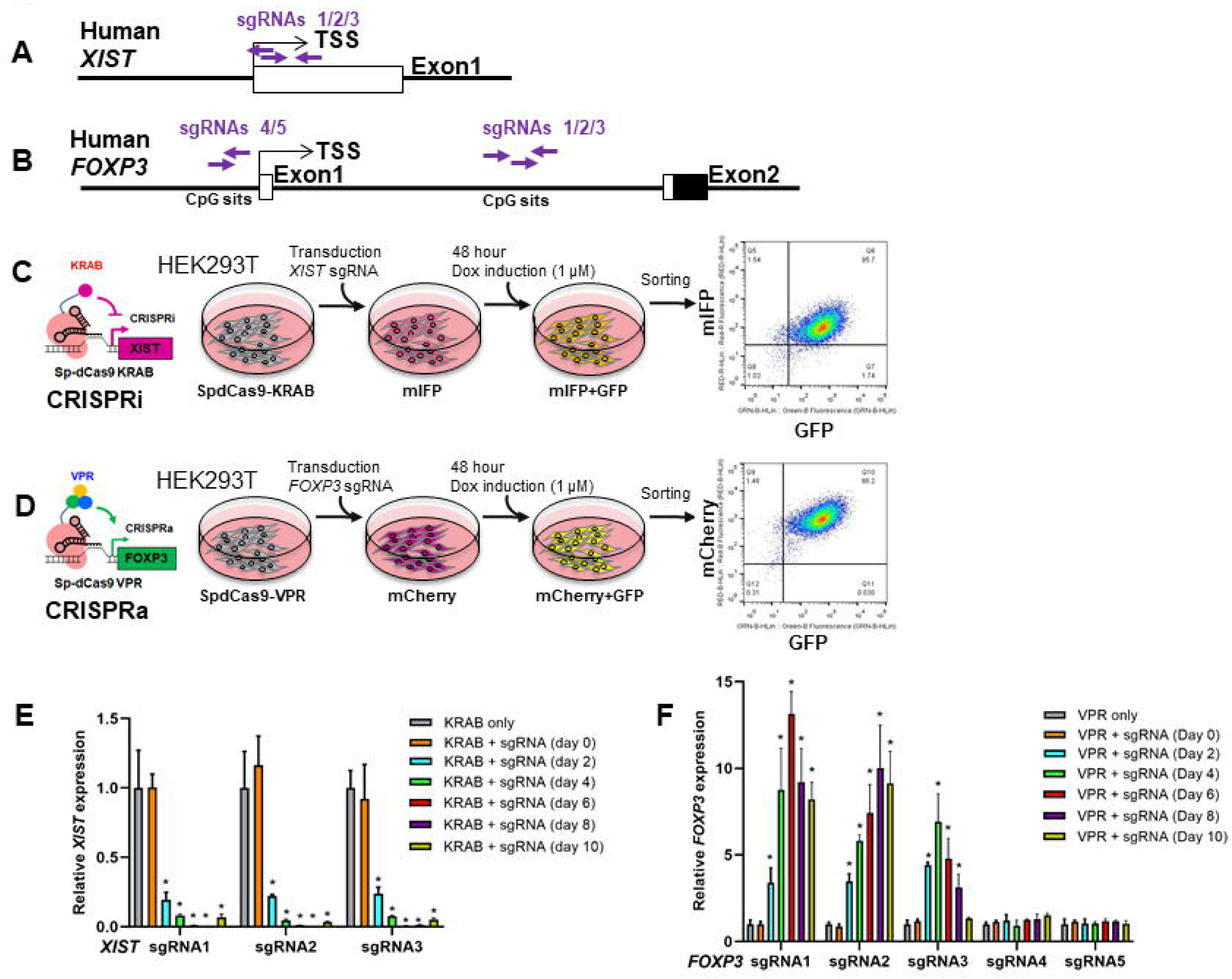
Targeted activation of *FOXP3* and inactivation of *XIST* in human embryonic kidney (HEK) 293T cells. **A,** sgRNAs 1/2/3 targeted to the -50 to +300 bp upstream of the transcription start site of the *XIST* locus for transcription repression. **B,** sgRNAs 1/2/3/4/5 targeted to the two CpG sites of the *FOXP3* proximal promoter and the intron 1 regions for transcription activation. **C,** CRISPRi experimental procedure for the transduction of *XIST* sgRNA (mIFP), Dox induction, and targeted cell sorting of SpdCas9-KRAB (GFP after Dox) stably expressing HEK293T cells. **D,** CRISPRa experimental procedure for the transduction of *FOXP3* sgRNA (mCherry), Dox induction, and targeted cell soring of SpdCas9-VPR (GFP after Dox) stably expressing HEK293T cells. **E, F,** quantitative expression analysis of *XIST* and *FOXP3* by qPCR of CRISPRi and CRISPRa cells, respectively. After Dox (1.0 μ g/ml) induction, the expression levels of *XIST* and *FOXP3* in the transduced cells were determined at days 0, 2, 4, 6, 8, and 10. The fold change in expression was calculated using the 2^-ΔΔCt^method with *GAPDH* mRNA as an internal control. Data are presented as the means ± standard deviation (SD). * *p* < 0.05 by ANOVA followed by Dunnet’s *post hoc* test *vs.* KRAB or VPR only group. CRISPRi, CRISPR interference; CRISPRa, CRISPR activation; sgRNA, single guide RNA; Dox, doxycycline; KRAB, transcription repressor Krüppel associated box for CRISPRi; VPR, transcription activators VP64-p65-Rta for CRISPRa. All experiments were repeated three times.

### Transcriptional regulation of *XIST* and *FOXP3* by CRISPRi/a in human HEK293T cells

In our previous study, we established two human embryonic kidney HEK293T stable cell lines derived from a female fetus that stably expressed SpdCas9-KRAB or SpdCas9-VPR (58) (Supplementary Table S3). First, we tested the repression of *XIST* in our CRISPRi system using the established SpdCas9-KRAB-stably expressing HEK293T cell line. Three sgRNAs were designed to target the promoter and enhancer loci of *XIST* (Fig. 1A). With CRISPRi HEK293T cells, we validated the repression of *XIST* transcription by the *XIST*-sgRNAs 1/2/3 (Figs. 1C and 1E) (46). Next, we tested the activation of *FOXP3* transcription in our CRISPRa system using the SpdCas9-VPR- stably expressing HEK293T cell line. Five sgRNAs were designed to target the promoter and enhancer loci of *FOXP3* (Fig. 1B). After addition of Dox, levels of the *FOXP3* transcript were increased by 3- to 13-fold by use of the *FOXP3*-targeting sgRNAs 1/2/3 in the CpG-rich site of the *FOXP3* intron 1, but not in the proximal promoter region of the transcription start site (Figs. 1D and 1F). This result validated the efficacy of our CRISPRi/a-based platform for targeted repression of *XIST* and activation of *FOXP3* in human HEK293T cells.

### Transcription regulation of *XIST and FOXP3* by CRISPRi/a in human breast cancer cells

Most human female breast cancer cell lines have low or no *FOXP3* expression (21), accompanied by heterozygous gene deletions but rare *FOXP3* mutations (Supplementary Table S4). Thus, in female breast cancer cells, the X-inactivated allele of *FOXP3* does not appear to undergo genetic changes during carcinogenesis, suggesting that it may be possible to reactivate *FOXP3* from XCI. We determined the expression levels of *XIST* in various female breast cancer cell lines, including MDA-MD-231, MCF7, and CRL2316, and compared levels to those in HEK293T cells. As shown in Supplementary Fig. S1, expression of *XIST* was high in CRL2316 and HEK293T cells, moderate in MCF7 cells, but low in MDA-MD-231 cells. In MDA-MD-231 cells, two alleles of *FOXP3* were no deletion and mutation but there were low expression levels of *FOXP3* (Supplementary Table S4), suggesting epigenetic inactivation at both active and inactive alleles of *FOXP3* in these cells.

To achieve simultaneous transcriptional repression of *XIST* and reactivation of *FOXP3* in the same cells, we utilized two orthogonal CRISPR/dCas9 systems (28), SpdCas9 and SadCas9. We used SpdCas9-KRAB for repression of *XIST* and SadCas9-VPR for reactivation of *FOXP3* (Figs. 2A-C). We chose MDA-MD-231 cells as a model to test the transcriptional regulation of *FOXP3* by CRISPRi/a, especially at active alleles. First, we established the CRISPRi/a MDA-MD-231 cell model stably expressing SadCas9-VPR and SpdCas9-KRAB (Fig. 2A and 2B and Supplementary Table S3). Then, we transiently co-transduced *XIST*- and *FOXP3*-sgRNAs into the CRISPRi/a MDA-MD-231 cells (Fig. 2C). The efficacy of transduction of sgRNAs in the cells was validated by fluorescence microscopy (Fig. 2D). Likewise, the expressions of SadCas9 and SpdCas9 were validated in the cells by Western blots (Fig. 2E). After transduction of sgRNAs, quantitative real-time PCR (qPCR) analysis showed that levels of the *FOXP3* transcript were increased 8-fold by the *FOXP3*-sgRNAs 1/2, with *XIST*- sgRNA 1 at day 4 and up to 8 days (Fig. 2F). However, levels of the *XIST* transcript appeared to be reduced by the *XIST*-sgRNAs 1/2 with *FOXP3*-sgRNA 1, after Dox addition, but this change was not statistically significant (Fig. 2G). Likewise, levels of the *FOXP3* transcript in MDA-MD-231 cells were not significantly changed after addition of Dox (Fig. 2F), suggesting a dominant activation of the *FOXP3* transcript from an active X-linked allele.

**Figure 2.**
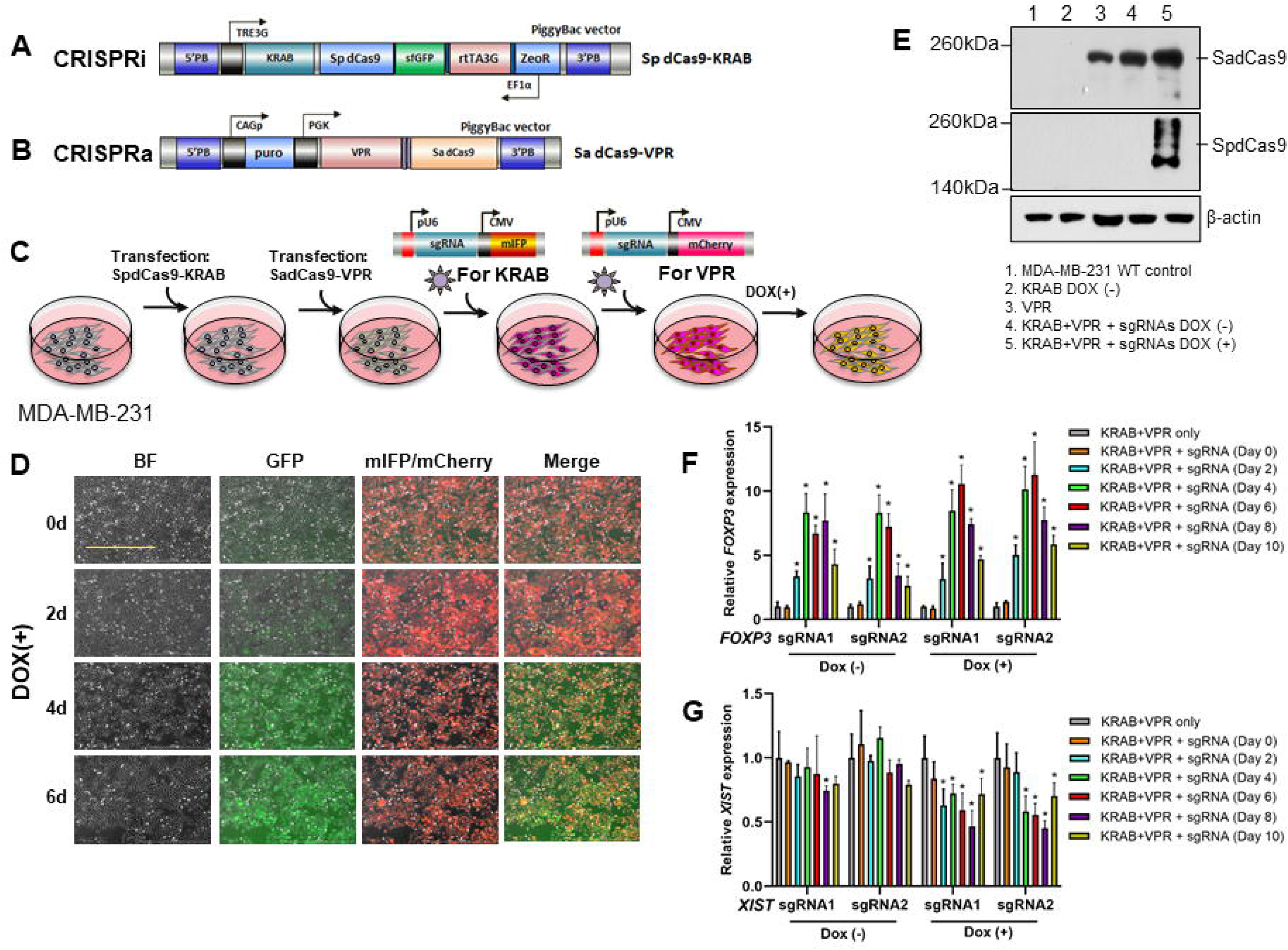
Assessment of CRISPRi/a in activation of endogenous *FOXP3* and repression of *XIST* in human breast cancer MDA-MB-231 cells. **A, B**, a diagram showing the constructs of CRISPRi/a, including *S. pyogenes* (Sp) dCas9-KRAB (SpdCas9-KRAB) and *S. aureus* (Sa) dCas9-VPR (SadCas9-VPR) used in the experiment. **C,** CRISPRi/a experimental procedure for the co-transduction of *XIST* (mIFP)- and *FOXP3* (mCherry)-sgRNAs, Dox induction, and targeted cell sorting of SpdCas9-KRAB (GFP after Dox) and SadCas9-VPR stably expressing MDA-MB-231 cells. **D,** efficacy of co-transduction of the *XIST* and *FOXP3* sgRNAs in CRISPRi/a cells before and after Dox induction at days 0, 2, 4 and 6 as determined by fluorescence microscopy. Scale bar, 1,000 μm. **E,** protein expression of SpdCas9 and SadCas9 in CRISPRi/a cells before and after sgRNA transduction and Dox treatment at days 0 and 4 as determined by Western blots with specific anti-SadCas9 and anti-SpCas9 antibodies. **F, G,** quantitative expression analysis of *FOXP3* and *XIST* before and after sgRNA transduction and Dox treatment of CRISPRi/a cells at days 0, 2, 4, 6, 8, and 10 as determined by qPCR. The fold change in expression was calculated using the 2^-ΔΔCt^ method with *GAPDH* mRNA as an internal control. Data are presented as the means ± SD. * *p* < 0.05 by ANOVA followed by Dunnet’s *post hoc* test *vs.* the KRAB+VPR-only group. All experiments were repeated three times.

In MCF7 cells, *FOXP3* has a heterozygous deletion (gene copy number, 0.67) with low levels of *FOXP3* expression (Supplementary Table S4). To validate the activation of the *FOXP3* transcript by CRISPRi/a, we established the CRISPRi/a MCF7 cell model and transiently co-transduced both *XIST*- and *FOXP3*-sgRNAs into these cells (Fig. 3A). The efficacy of sgRNA transduction was validated by fluorescence microscopy (Fig. 3B). After transduction of sgRNAs, levels of the *FOXP3* transcript were increased more than 8-fold by the *FOXP3*-sgRNAs 1/2, with *XIST*-sgRNA 1 (Fig. 3C). However, levels of the *FOXP3* transcript in the cells were not changed by Dox (Fig. 3C), although levels of the *XIST* transcript were reduced by Dox (Fig. 3D). These data support activation, in MCF7 cells, of the *FOXP3* transcript from an active X-linked allele.

**Figure 3.**
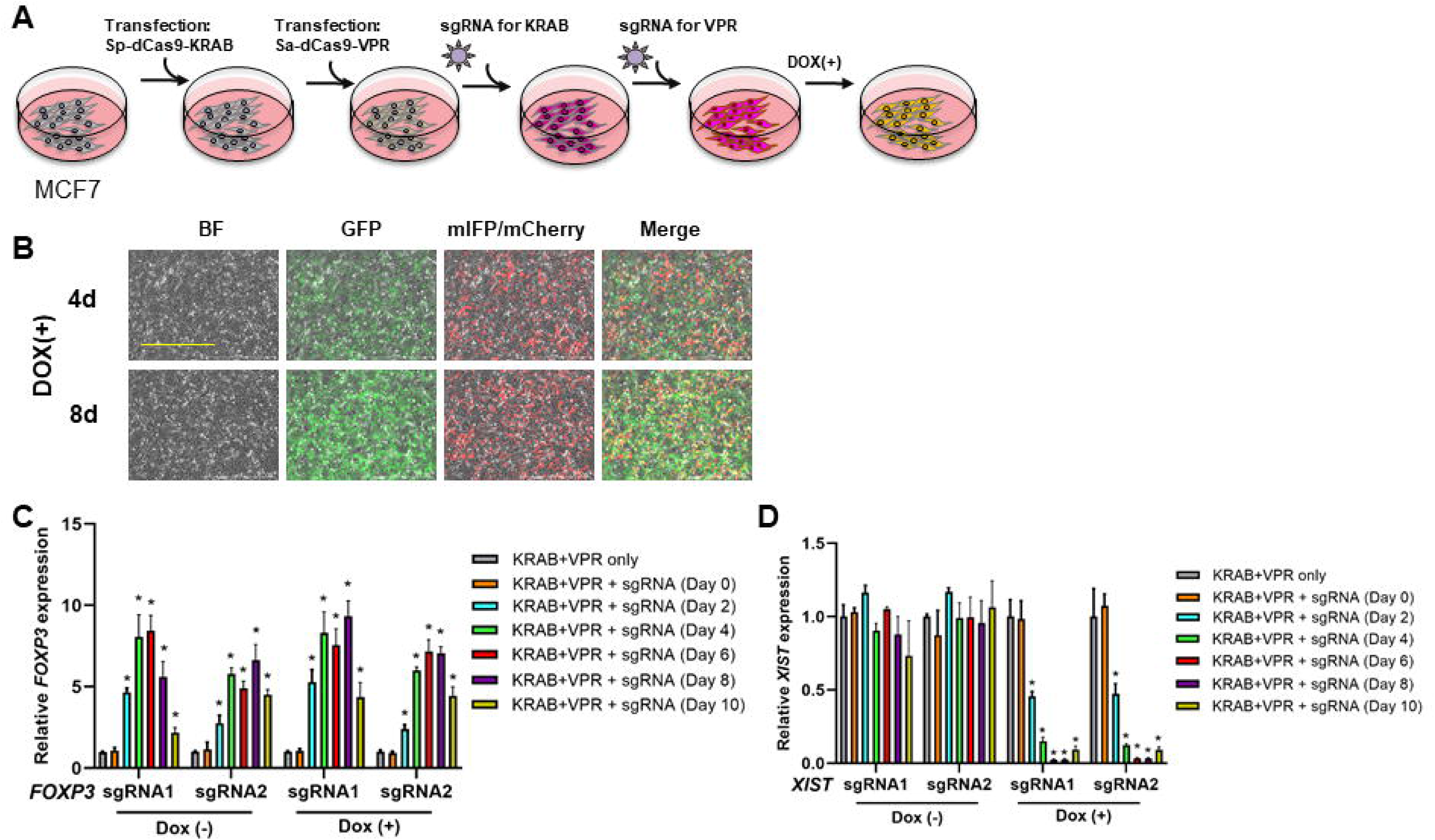
Assessment of CRISPRi/a to activate endogenous *FOXP3* and repress *XIST* in human breast cancer MCF7 cells. **A,** CRISPRi/a experimental procedure for the co-transduction of *XIST* (mIFP)- and *FOXP3* (mCherry)-sgRNAs and expression of SpdCas9-KRAB (GFP) by Dox induction in CRISPRi/a MCF7 cells. **B,** efficacy of co- transduction of the *XIST-* and *FOXP3-*sgRNAs in CRISPRi/a cells before and after Dox induction at days 4 and 8 as determined by fluorescence microscopy. Scale bar, 1,000 μm. **C, D,** quantitative expression analysis of *FOXP3* and *XIST* before and after sgRNA transduction and Dox induction of CRISPRi/a cells at days 0, 2, 4, 6, 8, and 10 as determined by qPCR. The fold change in expression was calculated using the 2^-ΔΔCt^ method with *GAPDH* mRNA as an internal control. Data are presented as the means ± SD. * *p* < 0.05 by ANOVA followed by Dunnet’s *post hoc* test *vs.* the KRAB+VPR-only group. All experiments were repeated three times.

### CRISPRi/a-mediated targeted reactivation of X-linked *FOXP3* from XCI in human breast cancer cells

We expect that, for human female breast cancer cells, our CRISPRi/a approach induces the transcription of *FOXP3* on both active and inactive X chromosomes. However, in MDA-MD-231 and MCF7 cells, our CRISPRi/a-mediated activation of endogenous *FOXP3* was most likely from the active X chromosome. The human breast cancer cell line CRL2316 (also known as HCC202), isolated from a female breast cancer patient, expresses a synonymous heterozygous mutation (p.L266L; c.798G>C) in the coding region of *FOXP3* (Supplementary Table S4), enabling us to determine, by cDNA sequencing, if *FOXP3* is reactivated from one or both alleles. Also, in CRL2316 cells, the two alleles of *FOXP3* showed no deletion but there were low expression levels of *FOXP3* (Supplementary Table S4). To test the CRISPRi/a-induced endogenous *FOXP3* from XCI, we selected CRL2316 as a cell model to assess the transcriptional regulation of *FOXP3* by CRISPRi/a at active and inactive alleles. First, we established the CRISPRi/a CRL2316 cell model (Fig. 4A and Supplementary Table S3). Then, we transiently co-transduced both *XIST*- and *FOXP3*-sgRNAs into these cells (Fig. 4A). After Dox induction, the sgRNA-transduced cells were sorted by flow cytometry (Fig. 4B). After sgRNA transductions, levels of the *FOXP3* transcript were increased approximately 6-fold by the *FOXP3*-sgRNAs 1/2 with *XIST*-sgRNA 1 (Fig. 4C). Transduction of *XIST* sgRNA reduced more than 90% of *XIST* expression at day 2 and up to day 10 after addition of Dox (Fig. 4D). Further, RNA fluorescence *in situ* hybridization (FISH) analysis showed a reduction of *XIST* at day 2, and most disappeared at day 4 (Fig. 4E) and up to day 10. As shown in Fig. 4E, at day 4 after Dox addition, the *XIST* probe was undetectable in most Dox-treated *XIST* sgRNA-transduced cells (bottom panels), as compared to Dox-untreated cells (top panels), suggesting a targeted inactivation of *XIST* on the inactive X chromosome by our CRISPRi approach. Simultaneously, levels of the *FOXP3* transcript were elevated approximately 2-fold at day 4 after Dox addition (Fig. 4C). Thus, for our established CRISPRi/a CRL2316 cells, we achieved the targeted repression and reactivation of endogenous *XIST* and *FOXP3*, respectively, together.

**Figure 4.**
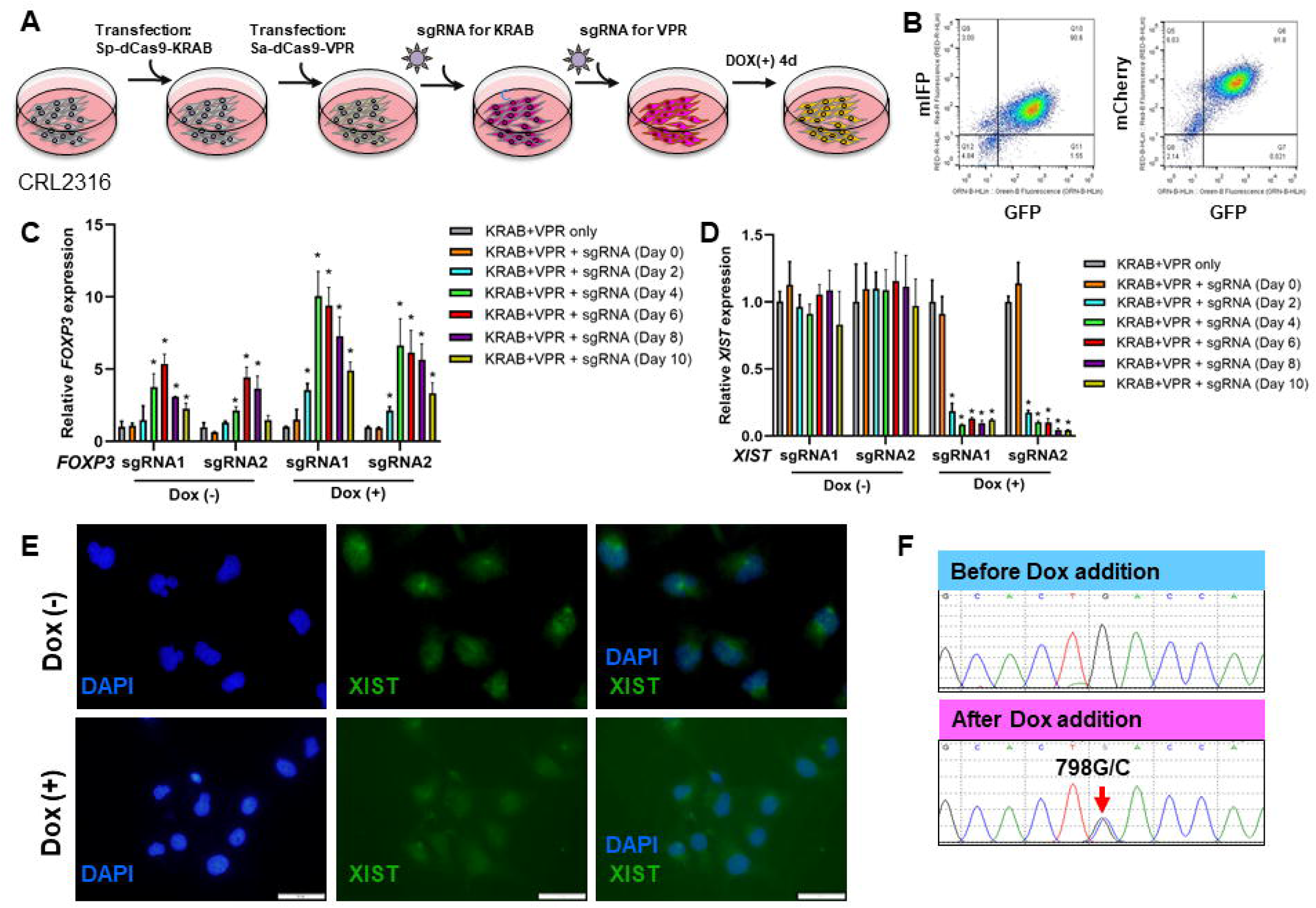
Assessment of CRISPRi/a to activate endogenous *FOXP3* and repress *XIST* in human breast cancer CRL2316 cells. **A,** CRISPRi/a experimental procedure for the co-transduction of *XIST* (mIFP)- and *FOXP3* (mCherry)-sgRNAs and Dox induction (GFP for SpdCas9-KRAB) in CRISPRi/a CRL2316 cells. **B,** targeted cell sorting of the CRISPRi/a cells after sgRNA transduction and Dox induction at day 4 as determined by flow cytometry. **C, D,** quantitative expression analysis of *FOXP3* and *XIST* before and after sgRNA transduction and Dox induction in CRISPRi/a cells at days 0, 2, 4, 6, 8, and 10 as determined by qPCR. The fold change in expression was calculated using the 2^-ΔΔCt^method with *GAPDH* mRNA as an internal control. Data are presented as the means ± SD. * *p* < 0.05 by ANOVA followed by Dunnet’s *post hoc* test *vs.* the KRAB+VPR-only group. **E,** expression of *XIST* in CRISPRi/a cells after Dox addition for 4 days as determined by FISH analysis. Cells were hybridized to the *XIST* probe (green). DAPI (blue) was used as a nuclear counterstain. Scale bar, 50 μ mutation analysis of a 798G/C mutation (red arrow) of the human *FOXP3* transcript in CRISPRi/a CRL2316 cells before and after activation of *FOXP3* as determined by cDNA sequencing. DAPI, 4’,6-diamidino-2-phenylindole. All experiments were repeated three times.

Next, we designed specific primers for mRNA surrounding the 798G/C mutant site of *FOXP3*. Total RNA was extracted from the *FOXP3*/*XIST* sgRNAs-transduced CRISPRi/a (CRISPRi/a-*FOXP3*/*XIST*) CRL2316 cells before and after Dox addition. As shown in Fig. 4F, by cDNA sequencing, a heterozygous 798G/C mutation of *FOXP3* was identified in Dox-treated CRISPRi/a-*FOXP3*/*XIST* CRL2316 cells but not in Dox-untreated CRISPRi/a-*FOXP3*/*XIST* CRL2316 cells, suggesting that CRISPRi/a-induced activation of the *FOXP3* transcript is at least partially reactivated from XCI under *XIST* downregulation.

Using the established CRISPRi/a CRL2316 cells, we determined the effect of CRISPRi/a-induced endogenous *FOXP3* on cell growth. We transiently transduced *XIST* sgRNA, *FOXP3* sgRNA, or both into CRISPRi/a CRL2316 cells for 48 hours, and then added Dox to the cells for 5 days. As shown in Supplementary Figs. S2A and S2B, levels of the *FOXP3* transcript were gradually elevated in the CRISPRi/a cells with *FOXP3* sgRNA, whereas levels of *XIST* were reduced in the CRISPRi/a cells with *XIST* sgRNA from days 1 to 5 after addition of Dox. Simultaneously, as shown in Supplementary Fig. S2C, cell growth was slower for CRISPRa cells with *FOXP3* sgRNA and slowest for CRISPRi/a cells with *FOXP3/XIST* sgRNAs relative to CRISPRi/a control cells, but there was no difference between cells with *XIST* sgRNA alone and control CRISPRi/a CRL2316 cells. These data suggest that CRISPRi/a-induced endogenous *FOXP3* inhibits growth of CRL2316 cells.

To exclude off-target effects of our CRISPRi/a, we evaluated the potential off-target genes of our designed sgRNAs using the online off-target searching tool (Cas-OFFinder, Daejeon, South Korea, http://www.rgenome.net/cas-offinder) (Supplementary Tables S5a-d). For *FOXP3*-sgRNA 1/2, the three nucleotide mismatched genes, *CFAP61*, *ERI3*, and *ZFAT*, were assessed by qPCR. Although expression levels of the potential off-target genes undulated in the CRISPRi/a cells with *FOXP3* sgRNAs, overall changes in these genes were not significant from days 0 to 10 after Dox addition (Supplementary Figs. S3A-C). Likewise, for *XIST*-sgRNA 1/2, the three nucleotide mismatched genes *ASXL2*, *IGF2BP2*, and *VAMP4*, were assessed by qPCR, but no overall change in these genes was evident from days 0 to 10 after Dox addition (Supplementary Figs. S3D-F).

### Effect of DNA demethylation on CRISPRi/a-mediated activation of *FOXP3* in human breast cancer cells

*XIST* RNA works in concert with DNA methylation and histone modifications to maintain XCI (14,59–62). Although reactivation of genes from XCI requires interference with *XIST* (20), *XIST* silence alone does not reactivate X-linked gene expression from XCI (14). In Tregs, DNA demethylation of the conserved CNS of the *FOXP3* intron 1 is specific for inducing or stabilizing transcription of *FOXP3* (47–50). Since the DNA methylation inhibitor 5-aza-2’-deoxycytidine (5-Aza-CdR) reactivates genes from XCI (63), we determined whether treatment with 5-Aza-CdR enhanced CRISPRi/a-mediated activation of *FOXP3* in human breast cancer cells. As shown in Fig. 5A, we transiently transduced *FOXP3*-sgRNAs into CRISPRa CRL2316 cells, followed by treatment with or without 5-Aza-CdR. The efficacy of *FOXP3*-sgRNA transduction was validated in the cells by fluorescence microscopy (Fig. 5B). The relative levels of the *FOXP3* transcript were quantified before and after treatment with 5-Aza-CdR during *FOXP3* activation. Although *FOXP3* was induced in the cells after *FOXP3*-sgRNA transduction, levels of the *FOXP3* transcript were not changed by treatment with 5-Aza-CdR (Fig. 5C). For the 10 CpG sites of conserved CNS of *FOXP3* intron 1, pyrosequencing analysis showed deregulation of DNA methylation by CRISPRa, 5-Aza-CdR, or both, in 7/10 CpG sites (Supplementary Fig. S4). However, these changes appeared to be not statistically significant (Figs. 6A-C). Next, we transiently co-transduced *FOXP3/XIST*-sgRNAs into CRISPRi/a CRL2316 cells, followed by treatment with or without 5-Aza-CdR and Dox addition (Fig. 5D). The expression of *FOXP3/XIST*-sgRNAs and Dox-induced SpdCas9-KRAB were validated in the cells by fluorescence microscopy (Fig. 5E). Treatment of *FOXP3*/*XIST*-sgRNAs-transduced CRISPRi/a cells with 5-Aza-CdR enhanced levels of the *FOXP3* transcript approximately 2-fold at days 2 and 4 (Fig. 5F), suggesting that induction of *FOXP3* by 5-Aza-CdR from XCI was under *XIST* downregulation. Likewise, pyrosequencing analysis revealed deregulation of DNA methylation by CRISPRi/a, 5- Aza-CdR, or both in most CpG sites (Figs. 6B and 6D and Supplementary Fig. S4).

**Figure 5.**
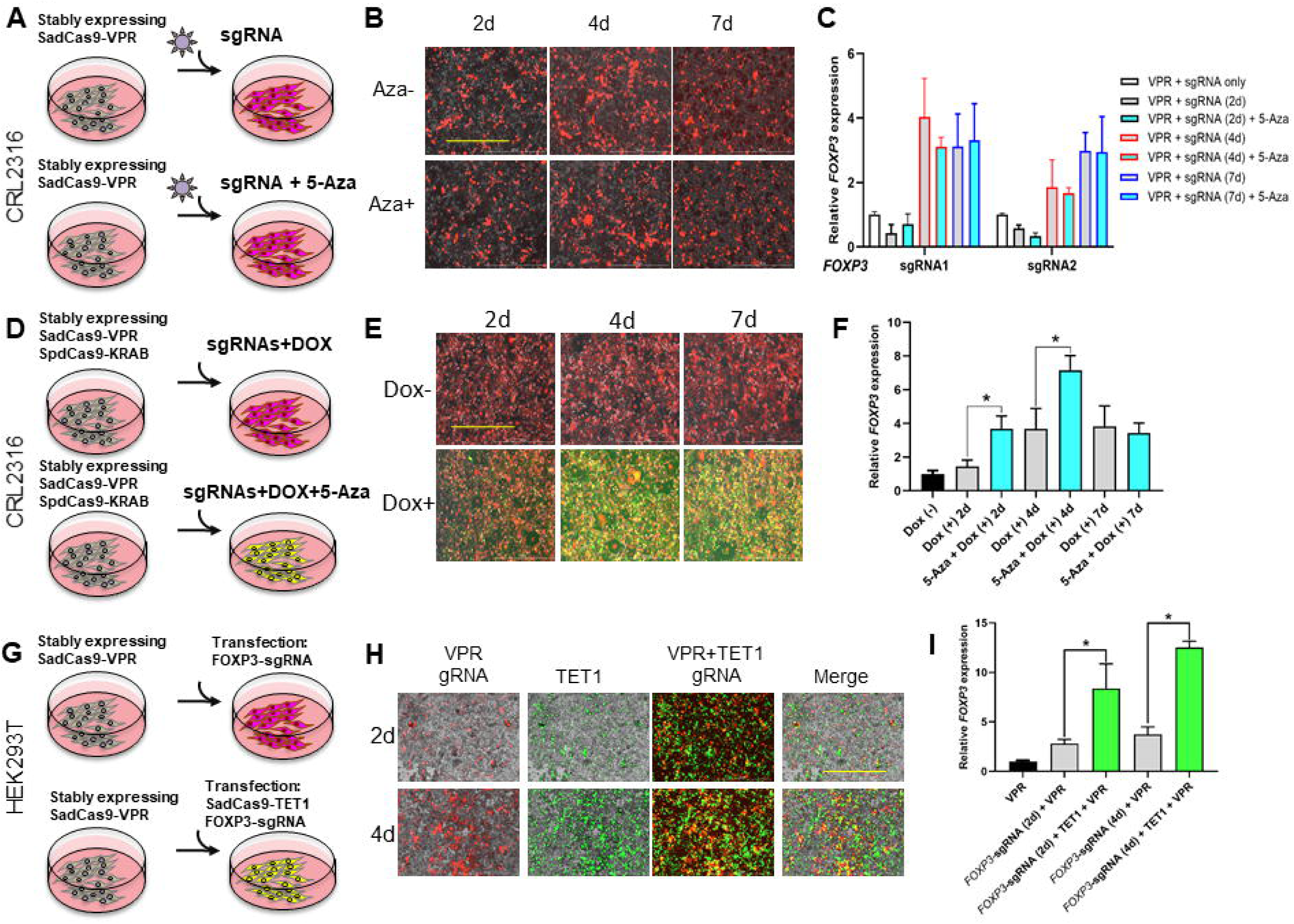
Effect of DNA demethylation on activation of endogenous *FOXP3* in human CRISPRi/a cells. **A,** the CRISPRi/a experimental procedure for the transduction of *FOXP3* sgRNA (mCherry) with or without 5-Aza-CdR (5-Aza) in CRISPRa CRL2316 cells. **B,** efficacy of transduction of *FOXP3* sgRNA in CRISPRa CRL2316 cells before and after 5-Aza treatment at days 2, 4, and 7 as determined by fluorescence microscopy. Scale bar, 1,000 μm **C,** quantitative expression analysis of *FOXP3* before and after sgRNA transduction and 5-Aza treatment of CRISPRi/a CRL2316 cells at days 0, 2, 4, and 7 as determined by qPCR. **D,** CRISPRi/a experimental procedure for the co-transduction of *XIST* (mIFP)- and *FOXP3* (mCherry)- sgRNAs with or without 5-Aza and Dox for CRISPRa CRL2316 cells. **E,** efficacy of co-transduction of the *FOXP3* sgRNA in CRISPRa CRL2316 cells before and after 5-Aza and Dox treatment at days 2, 4, and 7 as determined by fluorescence microscopy. Scale bar, 1,000 μm. **F,** quantitative expression analysis of *FOXP3* before and after sgRNAsc transduction and 5-Aza and Dox treatment of CRISPRi/a CRL2316 cells at days 0, 2, 4, and 7 as determined by qPCR. **G,** CRISPRi/a experimental procedure for the transduction of *FOXP3* sgRNA (mCherry) with or without SadCas9-TET1 (GFP) into CRISPRa HEK293T cells. **H,** efficacy of transduction of *FOXP3* sgRNA and transfection of SadCas9-TET1 into CRISPRa HEK293T cells at days 2 and 4 as determined by fluorescence microscopy. Scale bar, 1,000 μ m. **I,** quantitative expression analysis of *FOXP3* before and after sgRNA transduction and SadCas9-TET1 transfection into CRISPRi/a HEK293T cells at days 0, 2, and 4 as determined by qPCR. The fold change in expression was calculated using the 2^-ΔΔCt^method with *GAPDH* mRNA as an internal control. Data are presented as the means ± SD. * *p* < 0.05 by ANOVA followed by Dunnet’s *post hoc* test *vs.* a control group. All experiments were repeated three times.

**Figure 6.**
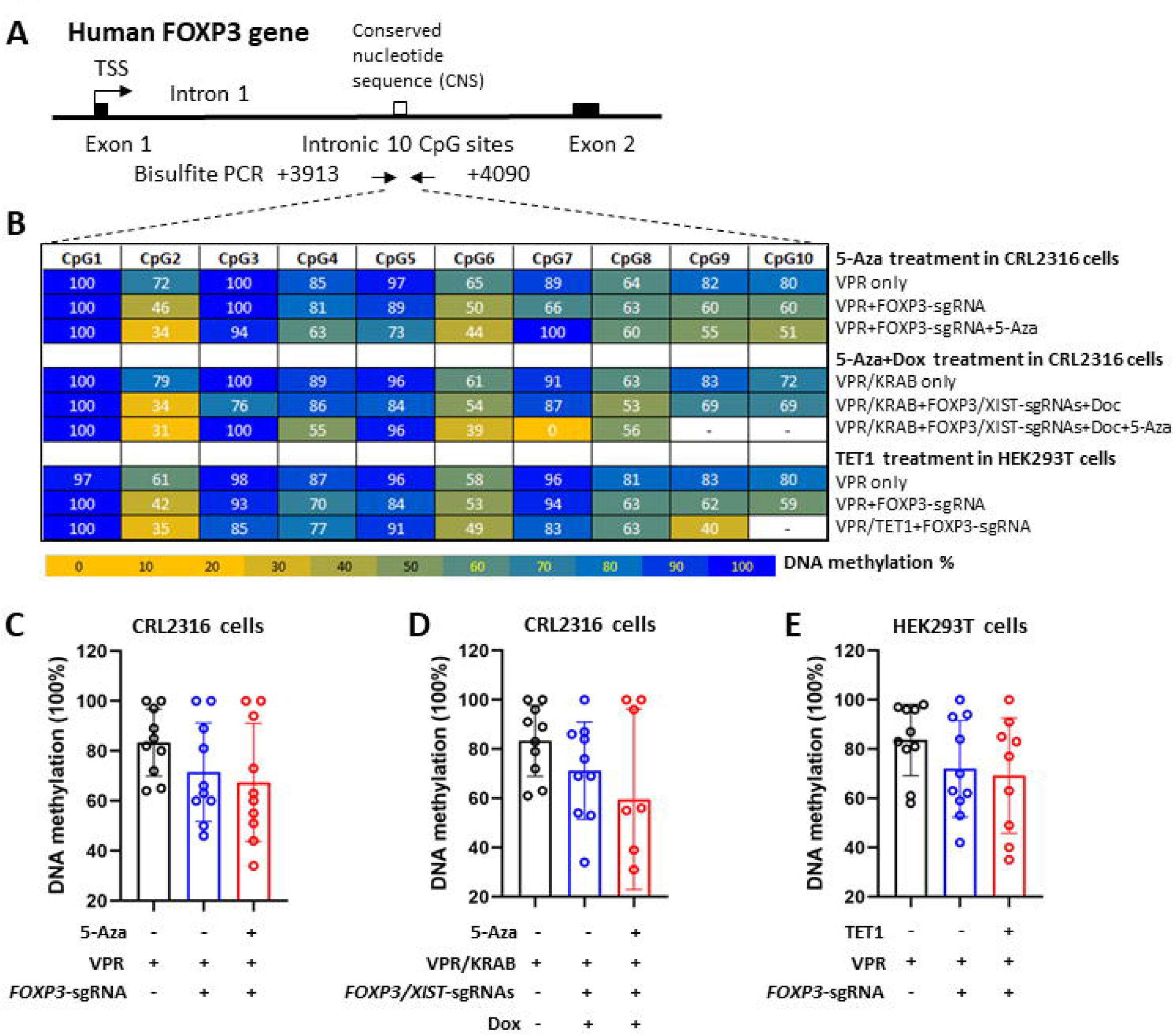
DNA methylation status of the rich CpG sites in the cconserved CNS of *FOXP3* intron 1 during CRISPRi/a-mediated activation of *FOXP3* in human CRL2316 and HEK293T cells. **A,** diagram of the 10 CpG sites in conserved CNS of *FOXP3* intron 1 and bisulfite PCR design for DNA methylation pyrosequencing analysis. **B,** heatmap of DNA methylation status determined by pyrosequencing. **C-E,** average levels of DNA methylation in CRISPRa CRL2316 cells, CRISPRi/a CRL2316 cells, and CRISPRa HEK293T cells before and after various treatments. Data are presented as the means ± SD. 5-Aza, 5-Aza-2’-deoxycytidine. All experiments were repeated three times.

The ten-eleven translocation (TET) family of DNA demethylase proteins converts cytosine methylated at C5 (5mC) to 5hmC, 5fC, and 5caC, and finally to cytosine with the aid of thymine-DNA glycosylase (64–70). These changes are associated with elevated gene transcription (71–73). Thus, we constructed SadCas9-TET1 (Supplementary Fig. S5A) for targeted DNA demethylation in the *FOXP3* CNS locus to enhance the activation of *FOXP3* in human breast cancer cells. The DNA construction was validated by gel electrophoresis analysis (Supplementary Fig. S5B). However, we failed to transfect SadCas9-TET1 into the CRISPRa CRL2316 cells due to the large construct size. Next, we transfected the SadCas9-TET1 and transduced the *FOXP3*-sgRNA into CRISPRa HEK293T cells (Fig. 5G). For these cells, the efficacies of transfection and transduction were validated by fluorescence microscopy (Fig. 5H). After 2 days of transfection, Western blots confirmed protein expression of SadCas9-TET1 in the transfected cells (Supplementary Fig. S5C). On days 2 and 4, levels of the *FOXP3* transcript were elevated approximately 3-fold in SadCas9-TET1-transfected and *FOXP3*-sgRNA transduced cells relative to cells transduced with *FOXP3*-sgRNA (Fig. 5I), suggesting, for HEK293T cells, synergistically enhanced activation of *FOXP3* by co-expression of TET1 and VPR. Likewise, pyrosequencing analyses validated, for most CpG sites, deregulation of DNA methylation by CRISPRa with VRP, TET1, or both (Figs. 6B and 6E and Supplementary Fig. S4).

### Histone modifications during CRISPRi/a-mediated activation of *FOXP3* in human breast cancer cells

Histone methylation and acetylation either repress or activate transcription (74). With the CRISPRi/a HEK293T cell model, we validated the expression of SadCas9-VPR protein by immunoprecipitation and Western blots with the SadCas9-specific antibody (Fig. 7A). To validate, in CRISPRi/a HEK293T cells, the specific binding of SadCas9- VPR to the intron 1 CNS locus of *FOXP3*, we performed chromatin immunoprecipitation (ChIP)-qPCR assays with a SadCas9-specific antibody with or without *FOXP3/XIST*-sgRNAs. As shown in Fig. 7B, binding of SadCas9-VPR to the intron 1 CNS locus of *FOXP3* was elevated more than 3-fold in cells transduced with sgRNAs relative to cells without sgRNAs; this binding was enhanced after addition of Dox to the cells. Although SadCas9-VPR also bound to *FOXP3* neighbor genes, *PPP1R3F* and *CCDC22*, these bindings were not elevated after transduction of *FOXP3/XIST*-sgRNAs or addition of Dox to the cells (Fig. 7C). Further, expressions of *PPP1R3F* and *CCDC22* in the cells were not changed after the transduction of *FOXP3/XIST*-sgRNAs and addition of Dox (Supplementary Figs. S6A and S6B). In addition, no specific binding of SadCas9-VPR was evident in the *XIST* locus (Fig. 7B). These data suggest a *FOXP3*-sgRNA-guided specific binding of SadCas9-VPR to the intron 1 CNS locus of *FOXP3*.

**Figure 7.**
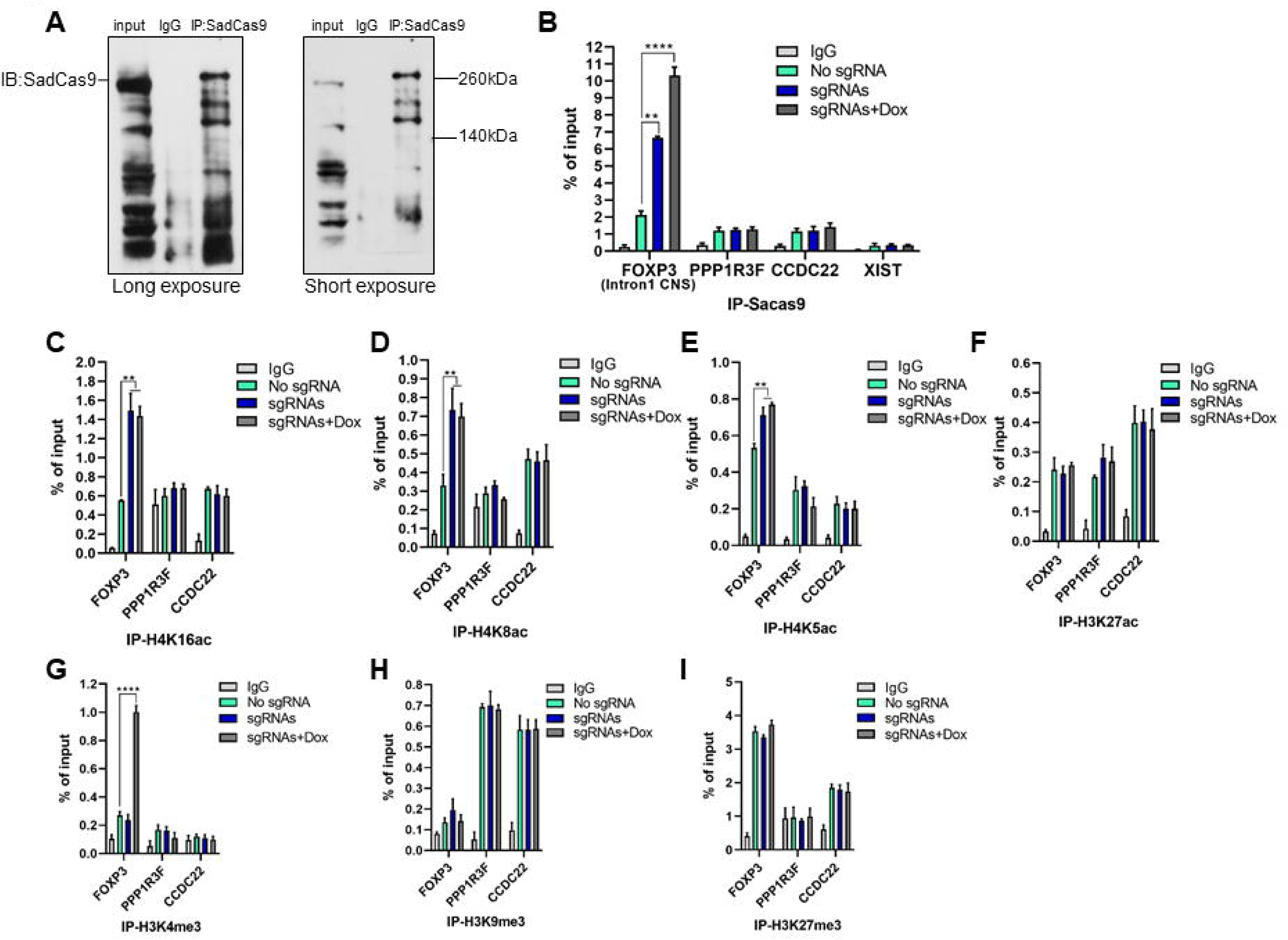
Histone modifications during activation of endogenous *FOXP3* in human CRISPRi/a HEK293T cells. **A,** immunoprecipitation (IP) with a specific anti-SadCas9 antibody (left panel: long exposure; right panel: short exposure) in CRISPRi/a HEK293T cells. **B,** SadCas9 chromatin-immunoprecipitation (ChIP)-qPCR in the conserved CNS of *FOXP3* intron 1 during CRISPRi/a-mediated activation of *FOXP3* with or without *XIST-* and *FOXP3*-sgRNAs and Dox at day 4 in HEK293T cells. The Y-axis represents % of input DNA. The promoter loci of *XIST* and *FOXP3* neighboring genes, *PPP1R3F* and *CCDC22,* were used as reference controls. **C,** global histone modifications during CRISPRi/a-mediated activation of *FOXP3* with or without *XIST-* and *FOXP3*-sgRNAs and Dox at day 4 as determined by Western blots of HEK293T cells. GAPDH was used as a loading control. **D-J,** various histone ChIP-qPCR analyses in the conserved CNS of *FOXP3* intron 1 during CRISPRi/a-mediated activation of *FOXP3* with or without *XIST-* and *FOXP3*-sgRNAs and Dox at day 4 in HEK293T cells. Data are presented as the means ± SD. ** *p* < 0.01 and **** *p* < 0.0001 by a two-sample *t*-test between two groups. All experiments were repeated three times.

To address the histone modification in the intron 1 CNS locus of *FOXP3* during activation of *FOXP3*, we performed a ChIP-qPCR assay with specific antibodies to H3K4me3, H3K9me3, H3K27me3, H3K27ac, H4K5ac, H4K8ac, and H4K16ac for CRISPRa-*FOXP3/XIST*-sgRNAs HEK293T cells. As shown in Figs. 7C-E, in CRISPRi/a cells after *FOXP3/XIST*-sgRNAs transduction and Dox addition, H4K5ac, H4K8ac, and H4K16ac were enriched in the intron 1 CNS locus of *FOXP3* but not in the promoter regions of *PPP1R3F* and *CCDC22.* Of note, during activation of *FOXP3*, H4K8ac and H4K16ac were elevated more than 2-fold in the intron 1 CNS locus (Figs. 7C and 7D), but there were no significant changes after Dox addition. However, in CRISPRi/a cells, H3K4me3, H3K9me3, H3K27me3, and H3K27ac were minimally changed in the *FOXP3*, *PPP1R3F*, and *CCDC22* loci after *FOXP3/XIST*-sgRNAs transduction (Fig. 7F-I). In addition, H3K4me3 was elevated more than 3-fold in the intron 1 CNS locus after addition of Dox (Fig. 7G). These data suggest that, during activation of FOXP3, *FOXP3*-sgRNA guided specific H4 acetylation at active alleles and H3K4 methylation at inactive alleles in the intron 1 CNS locus of *FOXP3*.

## Discussion

As reported here, we developed, for human female breast cancer cells, a CRISPRi/a approach for targeted transcriptional regulation of specific X-linked *FOXP3*, using two orthogonal dCas9-fusion systems, including SpdCas9-KRAB for CRISPRi to the *XIST* locus and SadCas9-VPR for CRISPRa to the *FOXP3* locus (Supplementary Fig. S7). We validated the efficacy of our CRISPRi/a system in inhibiting *XIST* transcription and activating *FOXP3* transcription simultaneously in these cells. The targeted reactivation of endogenous *FOXP3* from XCI was achieved by simultaneous use of CRISPRi/a. Of note, targeted reactivation of *FOXP3* inhibited growth of human female breast cancer cells. Furthermore, we optimized our CRISPRa system with the TET1 catalytic domain to enhance the transcriptional activation of *FOXP3*. The CRISPRi/a-mediated activation of *FOXP3* was accompanied by H4 acetylation at active alleles, including H4K5ac, H4K8ac, and H4K16ac, and H3K4 methylation at inactive alleles in the intron 1 CNS locus of *FOXP3*, indicating a CRISPRi/a-mediated epigenetic mechanism during activation of *FOXP3*.

Reactivation of a specific tumor suppressor gene with a targeted epigenetic modifier in cancer cells has been a goal of many cancer studies. Although CRISPRa by dCas9-VPR activates transcription of *FOXP3* in human female breast cancer cells, it remained unclear whether the activation of *FOXP3* was generated from the active, inactive, or both X chromosomes. In MDA-MB-231 cells, *FOXP3* is at a low expression level but was induced 8-fold by CRISPRa. However, *XIST* downregulation by CRISPRi did not appreciably enhance the CRISPRa-mediated activation of *FOXP3*. Since MDA-MB-231 cells express a deficient level of *XIST* (75) and may not have XCI, activation of *FOXP3* in these cells may be from the active X chromosome. In MCF7 cells, *FOXP3* was more than 8-fold induced by CRISPRa but was not increased after *XIST* downregulation by CRISPRi. Since MCF7 cells express a moderate level of *XIST* and a low expression level of *FOXP3* with heterozygous gene deletion, activation of *FOXP3* is most likely from the active X chromosome. In CRL2316 cells, there is a heterozygous, synonymous G/C mutation of *FOXP3* with moderate expression of *FOXP3*, but *XIST* expression is high. *FOXP3* was induced approximately 6-fold by CRISPRa and further enhanced nearly 2-fold after *XIST* downregulation by CRISPRi. Of note, cDNA sequencing showed that induced *FOXP3* was from the G allele before Dox addition but from both G/C alleles after *XIST* downregulation. Thus, for female breast cancer cells, this is the first targeted reactivation of endogenous *FOXP3* from both active and inactive X chromosomes through co-targeted epigenetic regulation of *XIST* and *FOXP3* by simultaneous CRISPRi/a.

In Tregs, DNA methylation controls X-linked FOXP3 gene expression (48). Treatment of Tregs with 5-Aza-CdR causes DNA hypomethylation in the CNS locus of *FOXP3*, inducing the expression of *FOXP3* (76). In CRISPRa CRL2316 cells, SadCas9 induces transcription of *FOXP3*, but 5-Aza-CdR treatment is unlikely to enhance the expression of *FOXP3.* Since, in mouse embryonic fibroblasts, 5-Aza-CdR in combination with genetic ablation of *Xist* synergistically causes X-reactivation from XCI (77), we treated CRISPRi/a CRL2316 cells with 5-Aza-CdR and Dox simultaneously and observed a synergistic activation of *FOXP3*, suggesting that 5-Aza-CdR enhances CRISPRi/a-induced *FOXP3* transcription from XCI under *XIST* downregulation. Furthermore, in Tregs, DNA demethylation in the CNS locus of *FOXP3* is required for inducing the expression of *FOXP3* (47–50). Targeted demethylation of the promoter or enhancer by dCas9-TET1 activates gene expression or promotes an active chromatin state (78, 79). We used SadCas9-VPR and SadCas9-TET1 to co-target the CNS locus of *FOXP3* in HEK293T cells. Although this co-targeting was not under *XIST* downregulation, SadCas9-VPR-mediated activation of *FOXP3* was enhanced by SadCas9-TET1, suggesting an *XIST*-independent reactivation of *FOXP3* by TET1. Likewise, co-targeting of dCas9-TET1 and dCas9-VP64 synergistically reactivates the X-linked *CDKL5* from XCI (25). Thus, *XIST* is likely to be dispensable during the reactivation of X-linked genes from XCI by TET1 with transactivators such as VPR or VP64.

dCas9-VPR is the fusion of dCas9 to a tripartite VP64-p65-Rta, which is more potent than dCas9-VP64 (80). It recruits various transcriptional factors (42) and endogenous active histone marks (e.g., H3K4me and H3K27ac) (81) at the enhancer and promoter regions (82). dCas9-KRAB induces heterochromatin formation and decreases chromatin accessibility through endogenous repressive histone marks (e.g., H3K9me3 and H3K27me3) at the enhancer and promoter regions (26,43,83,84). In the present study, our ChIP-qPCR analyses revealed CRISPRi/a-induced H4 acetylation, including H4K5ac, H4K8ac, and H4K16ac, and H3K4 methylation in the intron 1 CNS locus of *FOXP3*. H4K5ac facilitates the efficient deposition of the centromere-specific histone H3 variant CENP-A into centromeres (85), and H4K8ac is a potential regulator of chromatin-linked transcriptional changes (86). However, its role in cancer cells is still unknown. H4K16ac, which is associated with both transcriptional activation and repression (87), activates gene transcription by affecting both chromatin structure and interplay with non-histone proteins (88). Of note, a reduction of H4K16ac is evident in nearly all human tumors and cancer cell lines (89). Further, H4K16ac is enriched on the male X chromosome (90). Our data showed that, during activation of *FOXP3*, *FOXP3*-sgRNA-guided SadCas9-VPR but not *XIST*-sgRNA-guided SpdCas9-KRAB induced expression of H4K16ac in the intron 1 CNS locus. Thus, H4K16ac may be essential during the activation of *FOXP3* from the active X chromosome. In addition, H3K4me3 was elevated under *XIST* downregulation, suggesting that *FOXP3*-sgRNA guided specific H3K4 methylation in the intron 1 CNS locus of *FOXP3* may be from the inactive X chromosome. However, the epigenetic mechanism of the CRISPRi/a-regulated reactivation of X-linked *FOXP3* remains to be systematically investigated.

## Conclusions

The present study provides a better understanding of the CRISPRi/a-mediated activation of X-linked endogenous *FOXP3* and its regulatory mechanism in human female breast cancer cells. Also, our identification of the reactivation of the X-linked *FOXP3* from XCI moves beyond an incremental advance in breast cancer therapy by a targeted reactivation of X-linked tumor suppressor genes. Since epithelial *FOXP3* is inactivated in 70% of breast cancer samples (21), our results may lead to the design of preclinical studies to develop more effective treatments for female breast cancers with *FOXP3* dysfunction. In addition, this concept and tools may provide new routes of targeted therapy for other X-chromosome-linked genetic disorders.

## Materials and Methods

### CRISPRi/a DNA construction

Individual CRISPRi/a PiggyBac DNA constructs for dCas9 effectors used in this study are described in Supplementary Table S1. To assemble the doxycycline (Dox)-inducible dCas9-effector constructs, human codon-optimized *S. pyogenes* dCas9 was fused at the C-terminus with an HA tag and two SV40 nuclear localization signals, followed by the effector (26). The VPR effector (80) was assembled by fusing the activation domain of VP64 with the activation domain of p65 and Rta with two glycine-serine linkers. An extra SV40 nuclear localization signal was inserted between VP64 and p65. To pick out cell clones containing this vector, the puromycin-resistance gene was added to the N-terminal of the VPR effector followed by a CAG promoter. The *S. Aureus* dCas9 (gift from Feng Zhang, Addgene plasmid no. 61594) constructs and the VPR effector were driven by a PGK promoter. The catalytic domain of TET1 was amplified by PCR without any mutation and cloned onto the vector instead of the VPR effector. For visualization, enhanced green fluorescent protein (GFP) was fused to the N-terminal of the effector to replace the puromycin-resistance gene.

The *XIST*-sgRNAs and *FOXP3*-sgRNAs were designed to target the -50 to +300 bp upstream of the transcription start site of the *XIST* locus for KRAB transcription repression and target to the two CpG-rich sites of the promoter and intron 1 of the *FOXP3* locus for VPR transcription activation or TET1 DNA demethylation. As shown in Supplementary Table S1, for targeted repression of *XIST,* we constructed three *XIST*-sgRNAs to the pHR lentiviral U6-based expression vector (pSLQ2837), which expresses monomeric infrared fluorescent protein (mIFP). Likewise, we constructed five *FOXP3*-sgRNAs to the pHR lentiviral U6-based expression vector (pSLQ2806), which expressed monomeric red fluorescent protein (mCherry). Alternative sgRNA sequences (Supplementary Table S2) were generated by PCR reactions with Q5-High-Fidelity DNA-polymerase (NEB) according to the manufacturer’s protocol. We used pSLQ2837 (for *XIST* sgRNA)/pSLQ2806 (for *FOXP3* sgRNA) as PCR templates to the sgRNA using the forward primer and the universal reverse primer. Then, the purified PCR product was introduced by InFusion cloning into the pSLQ2837/pSLQ2806 backbone vector, which was digested with *BstX*I and *Xho*I as previously described (26). All DNA constructs of sgRNAs were confirmed by DNA Sanger sequencing.

### Cell culture and transfection

HEK293T, MCF-7, and MDA-MD-231 cell lines (ATCC) were cultured in high-glucose DMEM media supplemented with 10% fetal bovine serum (FBS, Thermo Fisher Scientific, Waltham, MA). The CRL2316 (also known as HCC202, ATCC) cell line was routinely grown in RPMI-1640 medium supplemented with 10% FBS. All cell lines were cultured at 37 °C with 5% CO_2_ for less than six months, were authenticated by examination of morphology and growth characteristics, and were confirmed to be mycoplasma-free. Short tandem-repeat analysis for DNA fingerprinting was also used to verify the cell lines. To achieve high transfection efficiencies, HEK293T, MCF-7, MDA-MD-231, and CRL2316 cell lines were stably generated with Piggybac-based dCas9-effector constructs and then followed by transduction with suitable sgRNA vectors. One day before transfection, the cultured cells were seeded in 6-well plates and were grown to about 80% confluency on the day of transfection. To integrate the Piggybac-based dCas9-effector constructs into the genome, the PiggyBac plasmid containing the dCas9-effector and Super PiggyBac transposase (Systems Biosciences, Palo Alto, CA) plasmid were co-transfected at a ratio of 2.5:1 using Lipofectamine 3000 (Thermo Fisher Scientific). After incubation with the transfection complexes for 48 hours, stably integrated cells were selected by Zeocin and GFP for CRISPRi and puromycin for the CRISPRa vector for 14 days. Then, the transfected cells were isolated as single clones and confirmed by Western blots.

To evaluate the effectiveness of the sgRNAs-induced dcas9 system, stably integrated cells containing the CRISPRi or CRISPRa vector were subjected to transient transduction of sgRNA vectors. Cells were harvested for RNA isolation at 2, 4, 6, 8, and 10 days post-transduction and analyzed by reverse transcriptase qPCR. To establish the cells containing CRISPRi/a vectors and sgRNA vectors, the most effective sgRNA vectors for CRISPRi/a were lentivirally transduced into the stably integrated cells containing the CRISPRi/a vector. For lentivirus generation, HEK293T cells were seeded into two 10-cm plates and were grown to about 80% confluency on the day of transduction. Then, individual lentiviral sgRNA vectors were co-transduced with two lentiviral packaging plasmids, psPAX2 and pMD2G (Addgene, Cambridge, MA) at a ratio of 2:2:1 using polyethylenimine (Polysciences, Warrington, PA). After 18 hours of transduction, the medium was exchanged with fresh medium, and, at 48 hours after replacing the medium, the packaged lentivirus supernatant was collected and passed through a 0.45-µm filter. Targeted cells were transduced with packaged sgRNA lentiviruses and were sorted, at 48 hours post-transduction, with mIFP or mCherry by use of a BD FACS Aria II flow cytometer (Thermo Fisher Scientific). Then, the positive mIFP or mCherry fluorescence cells containing CRISPRi/a vectors were cultured for four passages for use in subsequent experiments.

### qPCR

Total RNA was extracted from cells using the TRIzol Reagent (Thermo Fisher Scientific), and 1 µg of total RNA was reverse-transcribed with High-Capacity cDNA Reverse Transcription kits (Thermo Fisher Scientific). For each qPCR reaction, primers and 20 ng of cDNA were mixed in a 10-µl template and amplified using an SYBR Green Supermix kit (Thermo Fisher Scientific) with a Light Cycler 480 II instrument (Roche, Basel, Switzerland). The primer sequences are listed in Supplementary Table S2. The fold-increases of mRNA expression of the genes of interest were calculated using the 2^-ΔΔCt^ method with *GAPDH* mRNA as an internal control.

### Co-IP and Western blots

At about 90% confluency, cells on 10-cm plates were washed with cold PBS and lysed in ice-cold buffer (20 mM Tris–HCl, pH 8.0), 150 mM NaCl, 1 mM EDTA, 1% NP-40) supplemented with protease inhibitors (Sigma-Aldrich, St. Louis, MO) on ice for 10 min. After centrifuging of the preparations, the lysates were aliquoted into two tubes and incubated with either the designated antibody or an appropriate IgG control for 16 hours at 4°C. Then, the immune complexes were precipitated with Rec-protein G-Sepharose^TM^ (Thermo Fisher Scientific), and the proteins were eluted from the pelleted beads with 2x SDS sample buffer for subsequent Western blot analyses. For Western blotting, 30 µg of whole-cell lysates was separated on SDS-polyacrylamide gels and transferred to polyvinylidene difluoride membranes (Millipore, Burlington, MA). The membranes were incubated with appropriate primary antibodies, followed by either an anti-rabbit or anti-mouse IgG HRP-linked secondary antibody. The details of antibodies are listed in Supplementary Table S6. Then, after incubation of the membranes with chemiluminescence reagents, the HRP signal was evaluated by exposure of the membranes to X-ray films.

### Fluorescence-activated cell sorting (FACS)

For obtaining stably integrated cells with the CRISPRi vector, cells were gated for GFP-positive expression, and the population of interest was sorted and cultured in medium containing Zeocin (1 µg/ml). For sorting of cell pools with sgRNA vectors, cells were gated for mIFP- or mCherry-positive expression for the CRISPRi or CRISPRa system, and the population of the highest positive reporter fluorescent protein was collected and cultured for four passages for subsequent experiments. Each positive fluorescence population was gated, normalized to the un-transduced control cells by an BD FACS Aria II flow cytometer, and analyzed by FlowJo software (BD Biosciences, Ashland, OR).

### RNA FISH

RNA-FISH was performed as previously described (91, 92). Cells were seeded on glass coverslips in 3.5-cm plates for 2 days. After washing with Hank’s balanced salt solution (Thermo Fisher Scientific), coverslips were rinsed with Cytoskeletal buffer (CSK buffer) on ice. For extraction, coverslips were kept in CSK buffer with 0.5% Triton X-100 and 2 mM VRC (vanadyl ribonucleoside complex, Sigma-Aldrich) for 10 min on ice and were treated with 4% paraformaldehyde solution for 8 min of fixation. To detect human *XIST*, an appropriate DNA probe G1A (Addgene, plasmid no. 24690), encompassing approximately 10 kb of genomic DNA of the XIST gene, was processed with Digoxigenin-11-dUTP (Roche) and Nick-translation mix (Roche) (93). For hybridization, 50 ng of labeled probe was mixed with 10 μg of human Cot-1 DNA (Roche) and 10 μg each of salmon sperm DNA and *E. Coli* tRNA. The probe was air-dried and denatured on an 80°C heat block with 100% formamide. Then, coverslips were hybridized on the slides with 20 μ l of RNasin overnight. On the following day, the coverslips were rinsed in 2× saline-sodium citrate (SSC with 50% formamide) at 37°C, in 2×SSC at 37°C, in 1×SSC at room temperature on a shaker, and finally in 4×SSC at room temperature for 2 min to equilibrate cells before detection. Then, coverslips were incubated with a secondary antibody of anti-digoxigenin fluorescein (Roche) for 1 hour at 37°C and counterstained with 4’,6-diamidino-2-phenylindole (DAPI) for 1 min. Finally, coverslips were mounted using Vectashield (Vector Labs, Burlingame, CA) mounting media and imaged with a BX43 microscope (Olympus, Tokyo, Japan).

### Immunofluorescence imaging

Stably integrated cells containing CRISPRi or CRISPRa vector or both CRISPRi and CRISPRa vectors were seeded on 6-well plates for transduction with sgRNA vectors. At selected times, the positive reporter fluorescent signal of cells was imaged with a Lionheart FX Automated Microscope (BioTek, Winooski, VT). Selected positions of each image were acquired under a bright field, with GFP and Texas Red filter cubes, and merged for visualization of cells containing both fluorescent signals.

### DNA methylation analysis

DNA methylation status was determined by PCR analysis of bisulfite-modified genomic DNA using pyrosequencing (PyroMark Q96 ID; Qiagen, Germantown, MD) according to the manufacturer’s instructions. The primers were designed by use of the Pyromark Assay Design Software package (Qiagen), and 10 CpG sites of the *FOXP3* intron region were selected for quantitative DNA methylation analysis. These CpG sites were associated with activation of *FOXP3* by CRISPRi/a. DNA bisulfite conversion was performed by use of EpiTect Bisulfite Kits (Qiagen) followed by PCR amplification with primers. The pre-PCR products were used for bisulfite pyrosequencing to determine DNA methylation levels for each CpG site. The means of DNA methylation levels in the 10 CpG sites were calculated to obtain an average percentage of methylation. The primers used are described in the Supplementary Table 2.

### ChIP assays

ChIP assays were performed as previously described (94). Stable cells with CRISPRi and CRISPRa vectors were transduced with 12 µg of sgRNA vectors for CRISPRi and CRISPRa effectorsc on 10-cm plates using Lipofectamine 3000. After 96 hours of transduction, the cross-linked samples were sonicated in SDS Lysis Buffer (1% SDS, 10 mM EDTA, and 50 mM Tris-HCl, pH 8.1) using a Fisherbrand^TM^ Model 120 Sonic Dismembrator (Thermo Fisher Scientific) to 200-1,000bp fragments. Sheared samples were centrifuged for 10 min at 13,000 rpm and 4°C, and the pellets were diluted with ChIP dilution buffer (0.01% SDS; 1.1% Triton X-100; 1.2 mM EDTA; 16.7 mM Tris-HCl, pH 8.1; and 167 mM NaCl). ChIP enrichment of each sample was conducted by adding 2-4 μg of each antibody. Then, the immune complexes were pulled down with Rec-protein G-Sepharose^TM^ (Thermo Fisher Scientific), and the DNA was eluted from the pelleted protein G agarose/antibody/protein complex with elution buffer (1% SDS, 0.1 M NaHCO_3_). ChIP-qPCR was performed to quantify the amounts of immune-enriched DNA fragments. The primers and antibodies are shown in Supplementary Tables S2 and S6.

### CRISPR off-target analysis

To exclude the potential off-target effects of each sgRNA, we screened the sequences of *XIST* and *FOXP3* sgRNAs using the off-target searching tool (Cas-OFFinder, Daejeon, South Korea, http://www.rgenome.net/cas-offinder). Unbiased off-target screening was performed on the human reference genome GRCh38 for canonical SpCas9 PAM sites using 20-base-pair target sites without protospacer adjacent motif (PAM) sequences (95). The criteria for settings were confined to four or fewer mismatches with no DNA or RNA bulge. The total number and the possible binding loci of off-target genes of *XIST* and *FOXP3* sgRNAs are summarized in Supplementary Tables S5a-d. Subsequently, to assess the specificity of sgRNA-induced effects on target sites, qPCR was conducted to measure the mRNA expression of off-target genes (mismatches ≤ 3) of *XIST* and *FOXP3* sgRNAs and the nearest neighboring genes (*PPP1R3F* and *CCDC22*) to the *FOXP3* locus. Primers for qPCR and sgRNAs are shown in Supplementary Table S2.

### Statistical analyses

Differences in outcomes between two groups were compared by two-sided *t*-tests. Analysis of variance (ANOVA), one-and two-way, were used to test for overall differences, followed by a Dunnett *post hoc* test for differences between groups. All data were entered into an Access database using Excel (Microsoft 365 ProPlus) and analyzed with SPSS (version 25; IBM, Armonk, NY) and GraphPad (Prism 8, San Diego, CA).

## Abbreviations

5-Aza-CdR: 5-aza-2’-deoxycytidine
ANOVA: Analysis of variance
ChIP: chromatin immunoprecipitation
CNS: conserved non-coding sequence
Co-IP: co-immunoprecipitation
CRISPRa: CRISPR activation
CRISPRi: CRISPR interference
dCas9: endonuclease-deficient CRISPR/Cas9 protein
Dox: doxycycline
FACS: fluorescence-activated cell sorting
FISH: fluorescence *in situ* hybridization
GFP: fluorescent green protein
H3K4me3: tri-methylation at the 4th lysine residue of the histone H3 protein
H3K9me3: tri-methylation at the 9th lysine residue of the histone H3 protein
H3K27ac: acetylation at the 27th lysince residue of the histone H3 protein
H3K27me3: tri-methylation at the 27th lysine residue of the histone H3 protein
H4K5ac: acetylation at the 5th lysine residue of the histone H4 protein
H4K8ac: acetylation at the 8th lysine residue of the histone H4 protein
H4K16ac: acetylation at the 16th lysine residue of the histone H4 protein
KRAB: Krüppel-associated box
mCherry: monomeric red fluorescent proteins
mIFP: monomeric infrared fluorescent protein
PAM: protospacer adjacent motif
PRC1/2: polycomb repressive complex 1/2
qPCR: quantitative real-time PCR
Rta: Epstein–Barr virus R protein
Sa: *Staphylococcus aureus*
sgRNA: single guide RNA
Sp: *Streptococcus pyogenes*
TET: ten-eleven translocation
Tregs: regulatory T cells
VPR: tripartite VP64-p65-Rta proteins
XCI: X-chromosome inactivation
XIST: X-inactive specific transcript.

## Acknowledgements

We thank Dr. Donald Hill for editorial assistance in preparing this manuscript.

## Authors’ contributions

LW, RL and LSQ designed the research approach; XC, ZX, SW, XL and RL performed the experiments; XC, RL and LW analyzed data; XC, SB and LW performed statistical analyses; LZ, DZ and LSQ provided key resources; XC made a draft of the paper; XC, ESR, RL, LSQ and LW revised and edited the paper.

## Funding

This work was supported by grants from the DOD (W81XWH-17-1-0017 for L. Wang and W81XWH-17-1-0018 for L.S. Qi) and the Breast Cancer Research Foundation of Alabama (L. Wang).

## Availability of data and materials

Results are based, in part, upon data generated by Cas-OFFinder (http://www.rgenome.net/cas-offinder).

## Declarations

### Ethics approval and consent to participate

Not applicable.

### Consent for publication

All authors have agreed to publish this manuscript.

### Competing interests

The authors declare that they have no competing interests.

## Notes

### Competing Interest Statement

The authors have declared no competing interest.

