## Supplementary Figures 1-7 for "Targeted Reactivation of X-linked Endogenous FOXP3 Gene from X-chromosome Inactivation in Human Female Breast Cancer Cells"

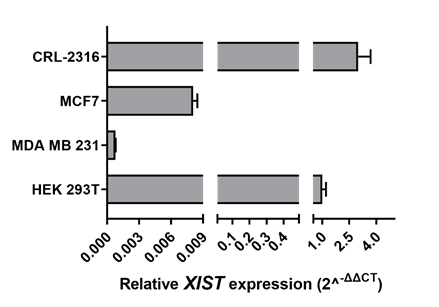

**Figure S1. Expression of *XIST* in human embryonic kidney (HEK) 239T cells and breast cancer cell lines.** The expression levels of *XIST* were assessed by qPCR. The fold change in expression was calculated using the 2^-ΔΔ Ct^ method with *GAPDH* mRNA as an internal control. Data are presented as means ± SD. HEK293T, a human embryonic kidney 293 cell line with the SV40 T-antigen; MCF7, a human estrogen receptor (ER)-positive breast cancer cell line; MDA-MB-231, a human triple-negative breast cancer (TNBC) cell line; CRL-2316, a human epidermal growth factor receptor 2 (HER2)-positive breast cancer cell line. All experiments were repeated three times.

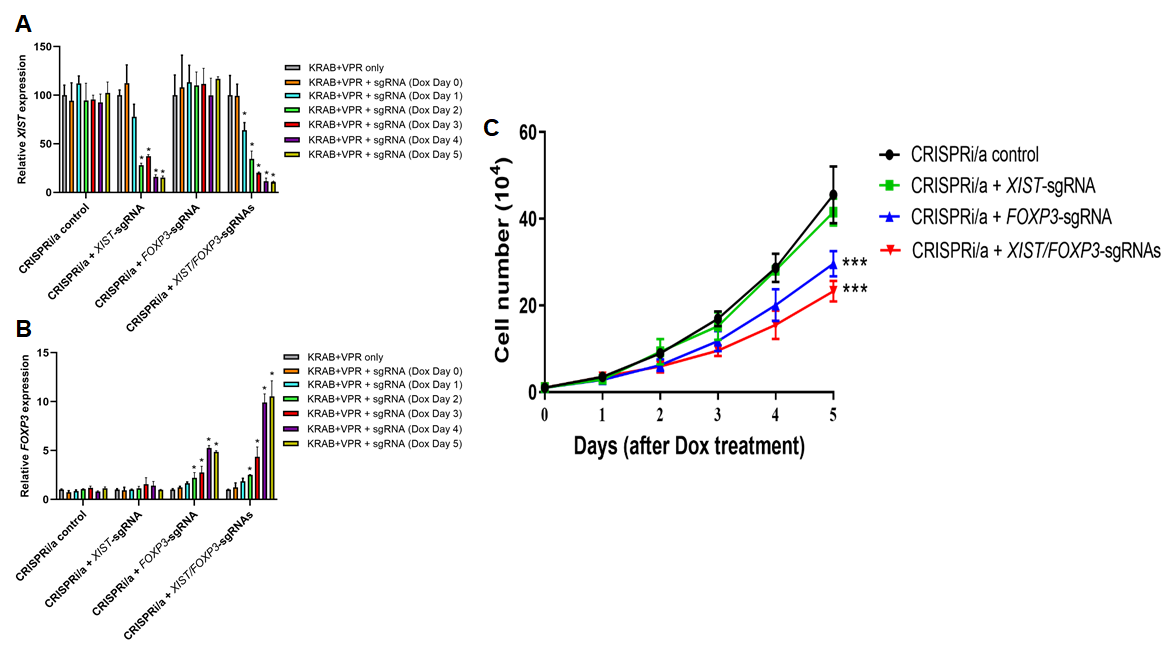

**Figure S2.** **The effect of targeted activation of *FOXP3* on cell growth in** **CRISPRi/a** **CRL2316 cells.** CRISPRi/a CRL2316 cells were transiently transduced with *XIST*-sgRNA, *FOXP3*-sgRNA, or both for 48 hours and then treated with Dox for 5 days. **A-B,** quantitative expression analysis of *FOXP3* and *XIST* by qPCR in cells with or without Dox. The fold change in expression was calculated using the 2^-ΔΔ Ct^ method with *GAPDH* mRNA as an internal control. Error bars, standard division. * *p* < 0.05 by one-way ANOVA followed by Dunnett’s analysis. **C,** effect of CRISPRi/a-induced endogenous *FOXP3* on growth of CRL2316 cells. Cell proliferation was measured after Dox treatment. Error bars, standard division. *** *p* <0.05 *vs*. CRISPRi/a control group by two-way ANOVA test. All experiments were repeated three times.

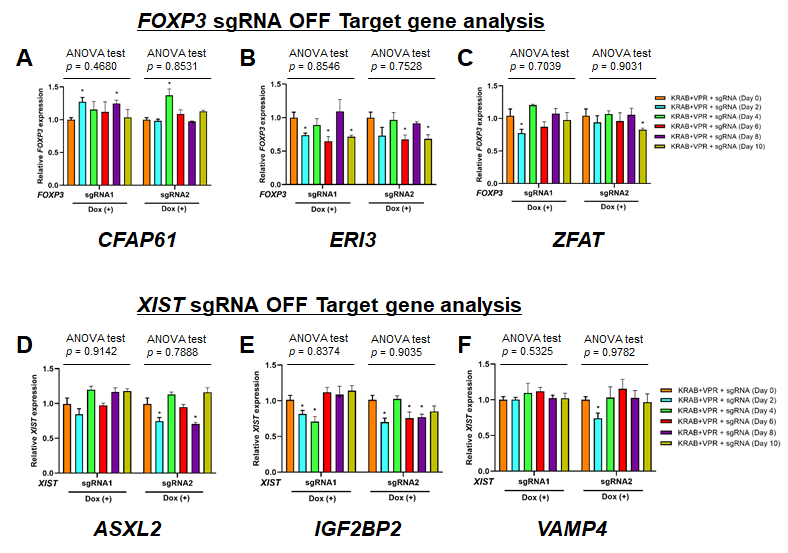

**Figure S3.** **Assessment of** **potential off-target** **genes of *FOXP3* and *XIST* sgRNAs in CRISPRi/a** **CRL2316 cells.** Quantitative expression analysis of the potential off-target genes of *FOXP3* sgRNAs (**A-C**) and *XIST* sgRNAs (**D-F**) before and after sgRNA transduction and Dox induction in CRISPRi/a CRL2316 cells at days 0, 2, 4, 6, 8, and 10 as determined by qPCR. The fold change in expression was calculated using the 2^-ΔΔ Ct^ method with *GAPDH* mRNA as an internal control. Data are presented as the means ± SD. * *p* < 0.05 by one-way ANOVA followed by Dunnett’s analysis. All experiments were repeated three times.

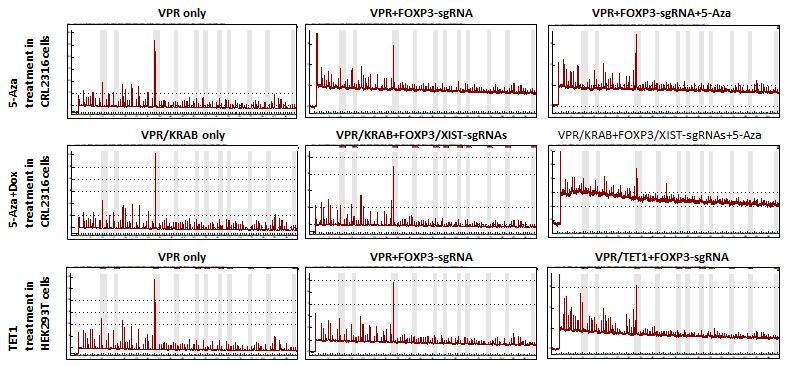

**Figure S4. DNA methylation analysis by pyrosequencing for CRISPRa CRL2316 cells, CRISPRi/a CRL2316 cells, and CRISPRa HEK293T cells with various treatments.** Pyrosequencing was performed to measure the methylation levels at 10 CpG sites in the conserved CNS of *FOXP3* intron 1 using the PyroMark Q96 ID pyrosequencer.

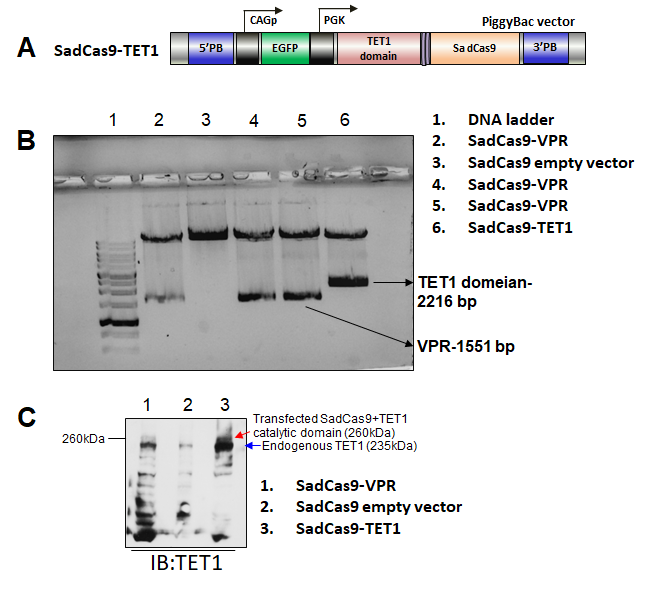

**Figure S5. Establishment of** **SadCas9-TET1 DNA constructs. A,** schematic construction of the SadCas9-TET1 vector used in the experiment. **B,** horizontal gel electrophoresis analysis of bands of the TET1 catalytic domain and VPR digested from SadCas9-TET1 and SadCas9-VPR vectors, respectively. Molecular sizes of the 10-kb DNA ladder are indicated on the left side. **C,** protein expression of TET1 after transfection into HEK293T cells. The SadCas9-TET1 vector was transiently transfected into HEK293T cells. The red arrow indicates the size of SadCas9-TET1 catalytic domain. The blue arrow indicates the full size of the endogenous TET1 protein. IB, Immunoblotting.

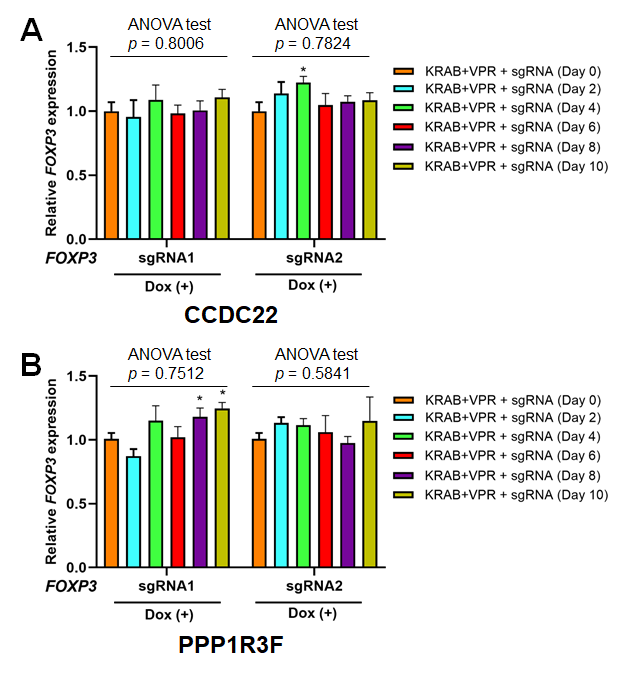

**Figure S6. Effect of CRISPRi/a-mediated activation of *FOXP3* on expression of its** **neighboring genes in CRL2316 cells.** *PPP1R3F* and *CCDC22* are two *FOXP3* neighboring genes at Xp11.23. Quantitative expression analysis of *CCDC22* (**A**) and *PPP1R3F* (**B**) before and after sgRNAs transduction and Dox induction in the CRISPRi/a cells at days 0, 2, 4, 6, 8, and 10 as determined by qPCR. The fold change in expression was calculated using the 2^-ΔΔ Ct^ method with *GAPDH* mRNA as an internal control. Data are presented as the means ± SD. * *p* < 0.05 by one-way ANOVA followed by Dunnett’s analysis. All experiments were repeated three times.

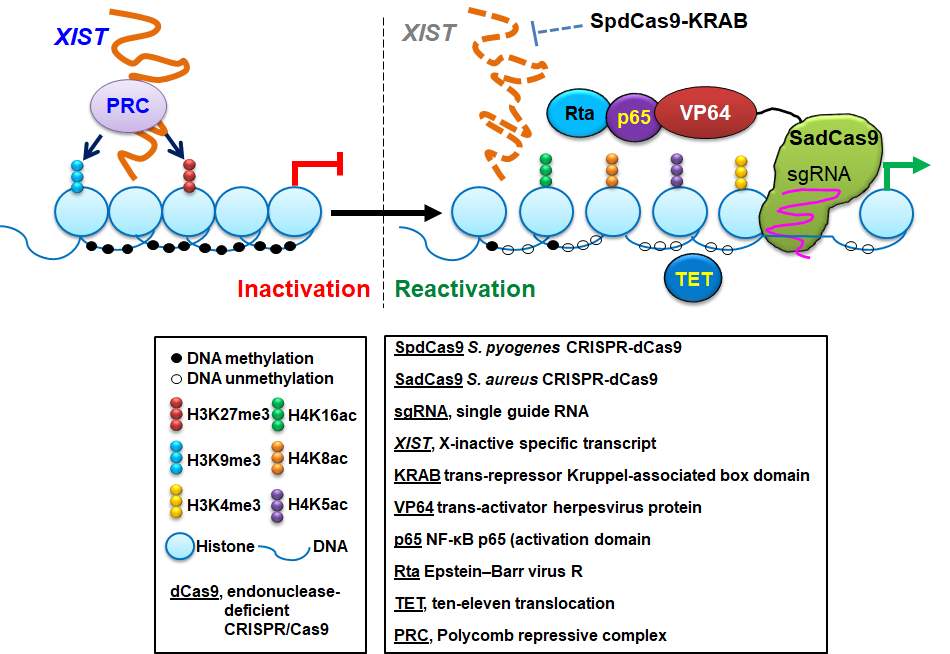

**Figure S7. Targeted reactivation of the X-linked endogenous FOXP3 gene from X chromosome inactivation (XCI) in female cells.** PRC (polycomb repressive complex) 1 or 2 recruits *XIST* RNA and promotes epigenetic modifications that block X-linked FOXP3 gene transcription on the inactive X chromosome. DNA binding by SadCas9-VPR (VP64/p65/Rta) and SadCas9-TET to the *FOXP3* intron 1 enhancer, and subsequent epigenetic modifications, in conjunction with the SpdCas9-KRAB to the *XIST* promoter, reactivates X-linked FOXP3 gene transcription from XCI.
